# Design principles for TB vaccines’ clinical trials based on spreading dynamics

**DOI:** 10.1101/249847

**Authors:** Sergio Arregui, Dessislava Marinova, Carlos Martín, Joaquín Sanz, Yamir Moreno

**Author notes:** Electronic address (Sergio Arregui). These authors contributed equally (Joaquín Sanz, Yamir Moreno).

## Abstract

Tuberculosis (TB) is one of the most complex diseases from the perspective of mathematical epidemiology. Individuals recently infected with the bacillus Mycobacterium tuberculosis can either develop TB directly in a matter of several weeks, or enter into an asymptomatic latent TB infection state (LTBI) that only occasionally derives into active disease, sometimes even decades after the infection event. The possible interruptions that a vaccine might provoke on these two mechanisms are indistinguishable in phase II clinical trials. In this work, we present a new methodology that allows differentiating vaccines that slow down the progression to disease from vaccines that prevent it. By introducing a stochastic framework for simulating synthetic clinical trials based on transmission models, we show how the method proposed here contributes both to reduce uncertainty in vaccine characterization and impact forecasts as well as to assist the design of clinical trials, improving their probabilities of success.

## I. INTRODUCTION

Despite the significant decline registered in TB burden during the last 20 years all over the world, Tuberculosis (TB) still remains one of the greatest menaces for public health world-wide. According to the Global TB Report of the World Health Organization (WHO) [1], 10.4 million people are estimated to have developed the disease during 2016 and 1.7 million people were killed by it. Furthermore, the prospects are not optimistic when we take into account the dramatic effects of HIV-co-infection [2, 3], with more than one million cases in 2016 [1], and the emergence of multidrug-resistent TB (MDR-TB) strains [4, 5], with almost 0.5 million cases in 2017 [1]. All these facts point to the indisputable need of new control methods and epidemiological interventions, and among them, the development of a new vaccine.

The role of the one-century old BCG vaccine at disrupting the TB transmission chain has been argued for decades [6–10]. As it turns out, BCG is unable to provide consistent efficacy levels when evaluated across different epidemic settings spanning different latitudes, individuals’ ages and levels of previous exposure to environmental Mycobacteria [10–12]. Consequently, among the different epidemiological interventions under current consideration, an improved vaccine holds the promise of a greatest impact in reducing TB cases and casualties; an observation that fuels the work of an active research community currently engaged in the development of disparate candidates for a new TB vaccine, 13 of which are nowadays being tested in clinical trials [13–15]. Since the resources available for this collective endeavor are limited, the development of rational approaches for filtering vaccine candidates represents a first order priority for Public Health Organizations and funding agencies. Accordingly, a well-defined stage-and-gate system has been defined and implemented, to which the different candidates must adhere in order to advance through the subsequent phases of the vaccine development pipeline. By providing a transparent framework for the evaluation and comparison of candidates at all stages, the system allows prioritizing the investment on the vaccines showing better performances regarding safety, immunogenicity and protection at each step, with the ultimate objective of maximizing the likelihood of success for the final finding of a new, safe and impactful TB vaccine.

Gathering the information needed to guarantee the eventual success of a vaccine poses a number of conceptual challenges that manifest at different stages of the clinical pipeline, making the evaluation and comparison of the different candidates under consideration an arduous task. The lack of protection correlates for TB [16, 17] hinders early efficacy evaluations, affecting our ability to identify the very presence of any protective effects from a given vaccine at the early stages of its development. That major limitation factually forces researches to rely new vaccines efficacy estimates to large randomized phase II and II-b clinical trials, which require the recruitment of thousands of individuals in high prevalence epidemiological settings during years to be completed. In this regard, the recent results from the nearly designed phase IIb trial of the novel vaccine MVA85A [18], even though it fails at providing evidence of significant protection, represents a solid framework for such studies.

However, even once the task of measuring the vaccine’s efficacy at reducing risk of disease (*V E_dis_*) is achieved upon completion of a phase II clinical trial, the interpretations of these results are not always immediate and can be due to different mechanisms of action of the vaccine. On the one hand, a vaccine can contribute to reduce disease risk by decreasing the fraction of fast progressors, increasing the probability that newly infected individuals are able to contain immediate bacterial proliferation developing latent TB infection (LTBI). Alternatively, the vaccine can delay the onset of active TB, slowing down the dynamics of fast progression instead of preventing it. These two possible mechanisms have different dynamical interpretations in terms of epidemiological transmission models, and might appear, in principle, independently and not necessarily correlated. The question of whether an individual will progress to active disease rapidly after infection or not depends on their ability of coordinating an initial strong innate immune response to the pathogen along with a later T-cell mediated adaptive response both of which are needed, if not to prevent infection by completely eliminating the bacteria, at least to arrest bacterial growth and confine the pathogen within granuloma [19]. If such complex response fails, fast transition to disease will take place, as a consequence of the host’s inability to both eliminate or confine bacteria around closed granuloma, yielding to rapid pathogen proliferation. Factors affecting the probability of individuals to join fast or slow paths to disease upon infection are both environmental and genetic, but little is known about how they impact, if they do it at all, the delay observed between infection and onset of symptoms in recent transmission TB cases (i.e. fast progressors) [19].

The main focus of this work is the analysis of this apparently innocent mechanistic degeneracy problem, in virtue of which, it is hard to distinguish, in the context of a clinical trial designed to estimate vaccine efficacy against disease, a vaccine that prevents from fast-progression from a vaccine that simply delays it. After a formal description of the question, we use Monte-Carlo methods to simulate synthetic clinical trials [20], along with a compartmental transmission model to produce impact forecasts [21] of vaccines that, yet compatible with single trial-derived lectures of *V E_dis_*, have different effects on fast-progression dynamics. Upon such exercise, we find that vaccines reducing the probability of fast progression are expected to offer, for the same observed levels of *V E_dis_*, significantly larger long-term impacts than vaccines that delay it. This situation translates into an excruciating level of uncertainty in what regards model-based impact evaluation of vaccines protecting against disease, unless a method for telling apart which of the two mechanisms constitutes a more plausible model under the light of the trial data is provided. Finally, we develop such a method for analysing raw results of clinical trials, which allows to isolate and measure what are the mechanisms of action of the vaccine that prevent or delay fast-progression to disease. This translates into more accurate impact forecasts and more faithful characterizations and comparisons of different types of vaccines, in a fraction of cases that depends on trials’ specifications (cohort sizes and follow-up periods duration).

## II. RESULTS

### A. Vaccine protection mechanisms against disease: reducing fast-progression probability vs. reducing fast transition rates to active TB

The essential goal of this work is to provide a means to interpret the results of a generic phase II clinical trial of vaccine efficacy in terms of compartmental models used downstream to evaluate vaccines’ impact and cost-effectiveness. In the more elementary version of these models (figure 1A) [5, 22–27], susceptible individuals get infected at a rate *β* defined by the interplay of their intrinsic susceptibility to infection and the frequency of contacts with infectious individuals that they experiment. Upon infection, they can develop fast-progression to disease (F) with a probability *p*, or slow-progression (latent infection L) with the remaining probability 1 − *p*. Finally, while fast progressors develop disease (D) at a rate *r* associated to typical transition times lower than one year, latent individuals can remain so for decades, only eventually falling sick, at a rate *r_L_*≪*r*. When we are talking about vaccinated individuals, parameters *β*, *p* and *r* get reduced to (1 − *ε_β_*)*β*, (1 − *ε_p_*)*p* and (1 − *ε_r_*)*r*, respectively.

**Figure 1.**
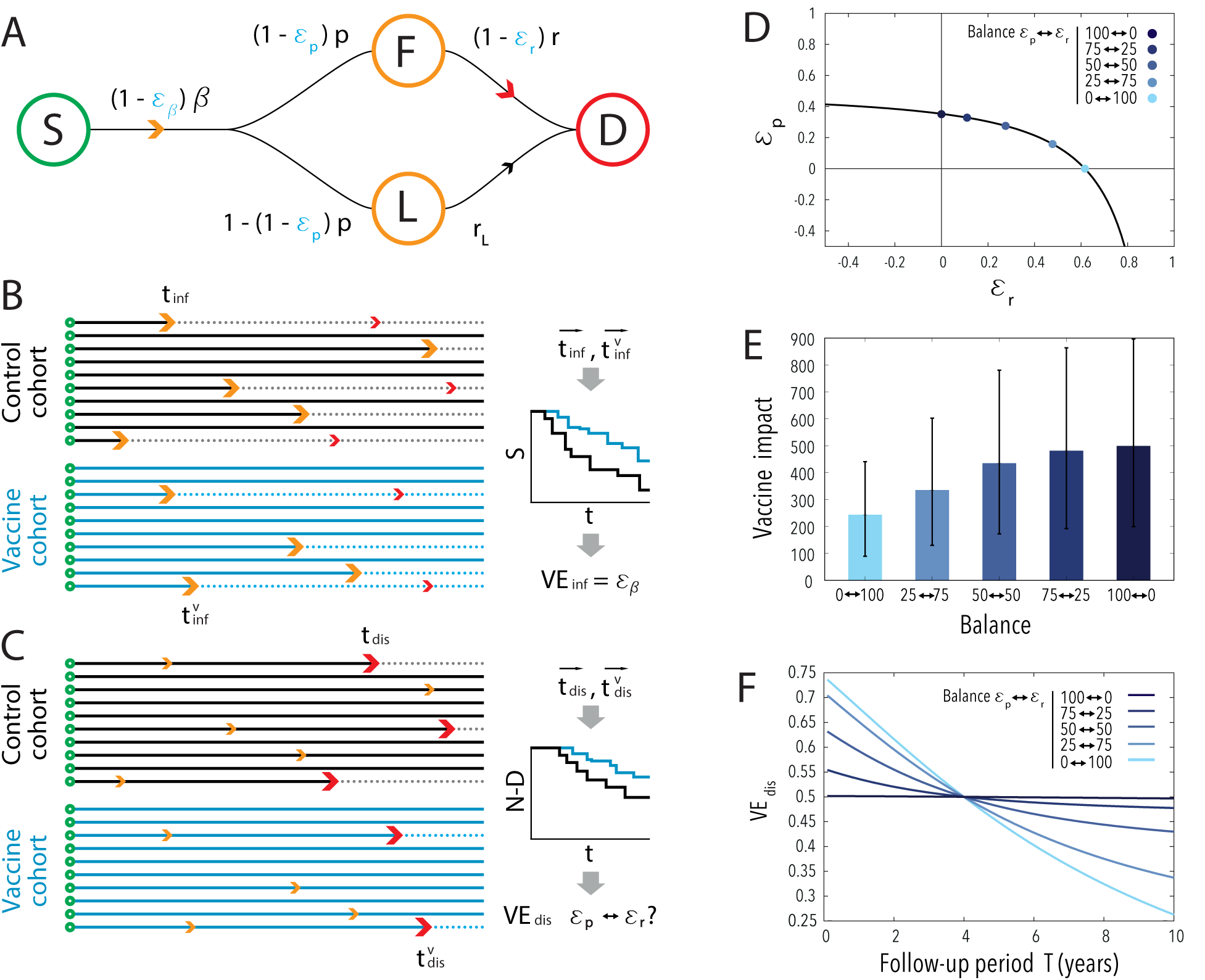
A: Transmission model. B: Scheme of times of transition to infection, and associated survival curves. C: Scheme of times of transition to disease (after infection) and associated survival curves (of non-disease states). D: Curve of values of (*ε_p_*,*ε_r_*) compatible with a measurement of *V E_dis_* = 0:5 after 4 years of follow-up (assuming *ε_β_* = 0). We have marked 5 different points in this curve, with very different balance between the two mechanisms that cause the degeneration, to be used in next examples. E: Forecasted impacts (thousands prevented cases) obtaining after introducing the 5 highlighted vaccines in a TB spreading model [21] in Ethiopia for the period 2025-2050. F: Evolution of measured *V E_dis_* for the 5 highlighted vaccines as a function of the follow-up period.

From this general picture, a series of assumptions are necessary to interpret the dynamics observed in a trial in terms of such minimal transmission model. A typical randomized, placebo controlled clinical trial of vaccine efficacy consists of two cohorts of initially susceptible individuals, (i.e. QuantiFERON-TB (QFT) negative [18]). At the beginning of the study, the vaccine is supplied to one of these cohorts, while a placebo is given to the other. Then, individuals will be periodically tested for infection (QFT) or disease (standard TB diagnosis criteria), during a given follow-up period short enough (i.e. less than 10 years) for us to assume that all the transitions to active disease observed were due to fast progressors in the first place. Finally, analyzing transition times to infection and disease end-points, two independent vaccine efficacy parameters can be measured: efficacy against infection *V E_inf_* and against disease *V E_dis_*, (figure 1, panels B,C). Under the light of this elementary transmission model, such efficacy observations can arise from the three putatively independent mechanisms: reduction of susceptibility to infection (via *ε_β_* > 0, see figure 1A), reduction of the fraction of fast-progressors (ε_p_ > 0) and reduction of the rate of fast progression to disease (*ε_r_* > 0).

All that said, the nature of the question under analysis turns evident: how to estimate three independent mechanisms of action of the vaccine from two measurements of vaccine efficacy? Regarding vaccine protection against infection, that question is easy to answer, for the only way for a vaccine to protect immunized individuals against infection is to reduce their probability of getting infected upon a contact with an infectious individual, which simply implies that *V E_inf_* = *ε_β_*. Instead, vaccine’s protection against disease offers a more complex outlook, for different vaccines, showing different combinations of effects on fast progression probabilities and transition rates to disease (*ε_p_* vs *ε_r_*) are compatible with a single lecture of *V E_dis_* (figure 1D). This implies that, from a trial-derived lecture of *V E_dis_* it is not possible to say whether the vaccine decreases individuals’ risk to develop disease by reducing the probability for them to become fast progressors upon infection or, alternatively, by slowing down the rate at which they develop TB.

Once the problem is identified, a direct way to quantify its relevance is to interrogate whether vaccines acting through different combinations of (ε_p_, ε_r_) that are still compatible with a common value of *V E_dis_* might produce different impacts when applied on large populations. To answer this question, we capitalized on a recently developed epidemiological model specifically designed to simulate different immunization campaigns of vaccines of different characteristics in different geographical settings (see Supplementary Appendix and [21]). Using such a model, we simulated the introduction of vaccines in Ethiopia, and obtained vaccines’ impact estimations in terms of TB cases prevented from 2025 to 2050, upon a newborns-focused vaccination campaign implemented at the beginning of that period (figure 1E, vaccines compatible with *V E_dis_* = 50% in a 4 year-trial). For this particular case, we found a difference of 256 thousand cases prevented (104-466 95% CI) between the first vaccine (ε_p_ = 0.00, ε_r_ = 0.74) and the last one (ε_p_ = 0.50, ε_r_ = 0.00), even though both vaccines provide an efficacy of *V E_dis_* = 50% in a 4 year-trial. This suppose a relative difference of 51% (45-59 95% CI)–, evidence the critical relevance of this degeneracy, which would compromise the reliability of any comparison between any disease-preventing vaccine candidates solely based on their observed efficacy levels *V E_dis_*.

These results have a straightforward interpretation: slowing down transition rates to disease of fast progressors will always be less effective, in the long term, than removing them from the fast-transition branch. A complementary way to illustrate this is to evaluate the time evolution of the efficacy *V E_dis_* that would be observed, for a single vaccine, in trials of different duration depending on its mechanisms of action (figure 1F), which can be derived analytically (see supplementary appendix, section S1). In this sense, a vaccine that reduces the intrinsic risk of developing fast progression (ε_p_ > 0) provides a reduction in the risk of disease that is almost independent of the observation time. Instead, for a vaccine that only slows down the rate at which fast-progressors develop active disease, the observed efficacy is a rapidly decreasing function of time: the longer the trial, the less efficient the vaccine would appear to be. This situation, which is intrinsically bound to the lower impacts forecasted from ε_r_-based vaccines, translates into a violation of the hypothesis of constant proportional risks underlying standard survival analysis (Cox regression) typically used to estimate *V E_dis_* for those vaccines, a question whose implications are discussed in detail in the supplementary appendix, section S2.

### B. Breaking the degeneracy: a method for estimating *ε_r_* and *ε_p_*

Even though violation of the proportional-hazard hypothesis in trials’ data constitutes a first signature suggesting delay in progression rates rather than fast progression prevention, its potential to tell apart quantitatively the possible mechanisms of action of a vaccine (see figure S1) is limited. Thus, we propose a different approach to provide an independent estimation of *ε_r_* and *ε_p_*. Our method consists of adding, to the custom estimation of V Einf and V Edis, a third statistical analysis aimed at directly estimating εr from the transition times between infection and disease (figure 2, panel A). To do so, unlike classical survival analysis, we only make use of data associated to not-censored times: that is, individuals that havecompleted their transition to disease within the follow-up period of the trial. By doing so,we can approximate that all these cases correspond exclusively to fast progressors (for *r_L_* ≪ *r*), and obtain an analytical expression for the expected distribution of transition times observed between infection and disease *t* = *t_dis_* − *t_inf_* (i.e., times between individuals first produce QFT positive results and they fall sick), conditioned to the moment the infection took place. This distribution *t* = Ψ (*r_cohort_*,*t_inf_*) has as its only parameter the transition rate of the cohort under analysis: either *r*, for the controls, or *r*(1 − ε*r*) for the vaccine group; parameters that we learn from the empirical data, across their respective uncertainties, using a maximum-likelihood approach (see Methods). By comparing the estimated transition rates from both cohorts, we get an estimation of *ε_r_*, and, from the analytical relationship *V E_dis_* = *f*(*ε_r_*; *ε_p_*), we usethe parallel estimation of *V E*_dis_ to easily resolve *ε_p_*, as detailed in the appendix (sectionS1). As a result, we solve the degeneracy issue, obtaining a full description of the vaccinethrough the individual estimations of the three effects (*ε_β_*, *ε_r_*, *ε_p_*).

**Figure 2.**
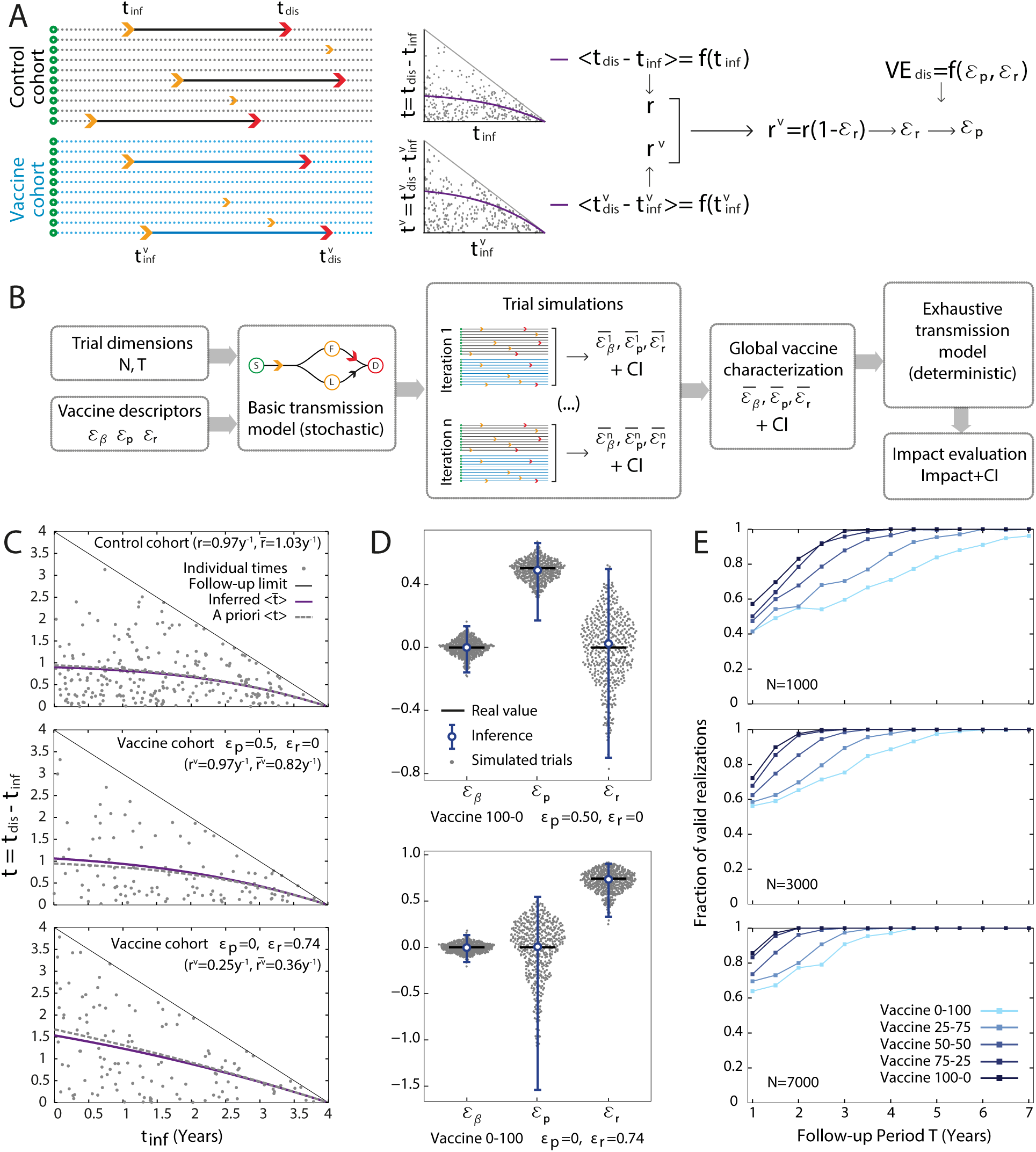
A. Scheme of the Method for characterizing vaccine’s efficacies. From the distribution of times of disease and infection at both cohorts we obtain *ε_r_*, and then *ε_p_* from the proportion of individuals that have suffered TB during the trial. B. Schematic representation of the process of this work. From a given vaccine (*ε_β_*; *ε_r_*; *ε_p_*) and trial design (*N*, *T*) we simulate 500 equivalent trials that we characterize using our method. From the inferred parameters we obtain a global estimation of the parameters and their CIs that we will use to evaluate their impact and their respective uncertainty. C. Transition times of control cohort (top) and vaccinated cohorts of an *ε_p_*-based vaccine (center) and an *ε_r_*-based vaccine (bottom). Transition times from infection to disease are considered to follow an exponential distribution truncated by the finite follow-up period of the trial (black line), whose expectation values (blue for vaccinated cohorts, grey for controls) and residual variances depend on time at infection as a consequence of the truncation. Using likelihood maximization, we infer within-cohort transmission rates to disease which generate expectation values curves for the transition times that closely resembles the a priori known analytical predictions for medium-size trials (dashed lines). D. Probability density of the inferred parameters of the vaccines (*ε_β_*; *ε_r_*; *ε_p_*), alongside the inferred parameters (with their respective CIs) for two different vaccines: *ε_p_*-based (top) and *ε_r_*-based (bottom). E: Fraction of correct realizations of a trial (i.e. realizations from which we obtain a biologically possible characterization) as a function of the follow-up period, for three different cohort sizes and 5 different vaccines (same 5 vaccines remarked in Figure 1)

To test the performance of our approach, we used Monte Carlo methods to simulate synthetic clinical trials of different lengths and sizes, for vaccines acting through different mechanisms and showing disparate levels of protection (Figure 2, panel B). For a given cohort size *N* and follow-up period *T*, we use as inputs the a prioriknown three vaccine descriptors (*ε_β_*, *ε_r_*, *ε_p_*) to simulate, using a stochastic implementation of the elementary transmission model represented in figure 1A, the development of a possible realization of the trial. Thus, we generate a set of vectors of transition times to infection and active TB that summarizes it. Since the model is probabilistic by construction, we iterate to obtain a set of simulated trials describing a distribution of most-likely outcomes associated to such a trial set-up as a function of the a-priori known vaccine behavior.

Then, for each clinical trial realization, we use our method to infer the vaccine descriptors (*ε_β_*, *ε_r_*, *ε_p_*). In figure 2, panel C we represent three examples of how the transition times associated to a single realization of a synthetic trial can be used to infer the transmission rates at the control cohort (upper panel) or at the vaccine cohort used, which are compared to infer *ε_r_*. Here, two types of vaccines are considered: an example of a vaccine that prevents fast progression (center, where the inferred transition rates are not significantly different from the controls), and another example where the vaccine reduces transmission times to disease within the fast progression branch (lower panel). Then, once *ε_r_* is estimated from the analysis of the transition times, we infer the other two parameters as described above to retrieve estimates for the three vaccine descriptors (*ε_β_*, *ε_r_*, *ε_p_*). As it can be seen in Figure 2D, this closely reproduce the initial values used to simulate the trials. Results correspond to a set of 500 realizations for an example vaccine with *ε_β_* = 0, *ε_p_* = 0.4, *ε_r_* = 0.4.

A final question to address concerns the range of applicability of our method. Since the key step of the proposed approach relies on the calibration of the distribution of infection-to-disease progression times, the size and duration of it must be big enough to ensure sufficient statistics. In that sense, we define biologically meaningful intervals for the vaccine parameters *ε_r_* and *ε_p_*, and reject all the trial simulations that, due to insufficient statistics, derive into inferred parameters values that go beyond those intervals (see Methods section). In figure 2E we represent the fraction of stochastic realizations that yield valid inferences of vaccine descriptors, for the same five vaccines represented in figure 1E. For atrial with a cohort size of 3000 individuals and follow-up period of 4 years, only a vaccine acting exclusively through *ε_r_* yields a probability of observing a trial incompatible with our method that surpasses 10%.

### C. Method’s evaluation: uncertainty reduction in impact evaluations

In the previous section, we described how we simulated stochastic realizations of clinical trials of different vaccines, and inferred, from each realization, the vaccine descriptors (*ε_β_*, *ε_r_*, *ε_p_*), in a way that is blind to the “real” values used to generate the synthetic trials. Next, we are interested in quantifying the extent to which impact forecasts might benefit from having access to the specific values (*ε_β_*, *ε_r_*, *ε_p_*). To this end, we feed our epidemic model [21] with these parameters, simulating the introduction of the vaccines in Ethiopia in 2025, and estimate impact forecasts in number of TB cases prevented within the period 2025-2050 (Figure 3A).

**Figure 3.**
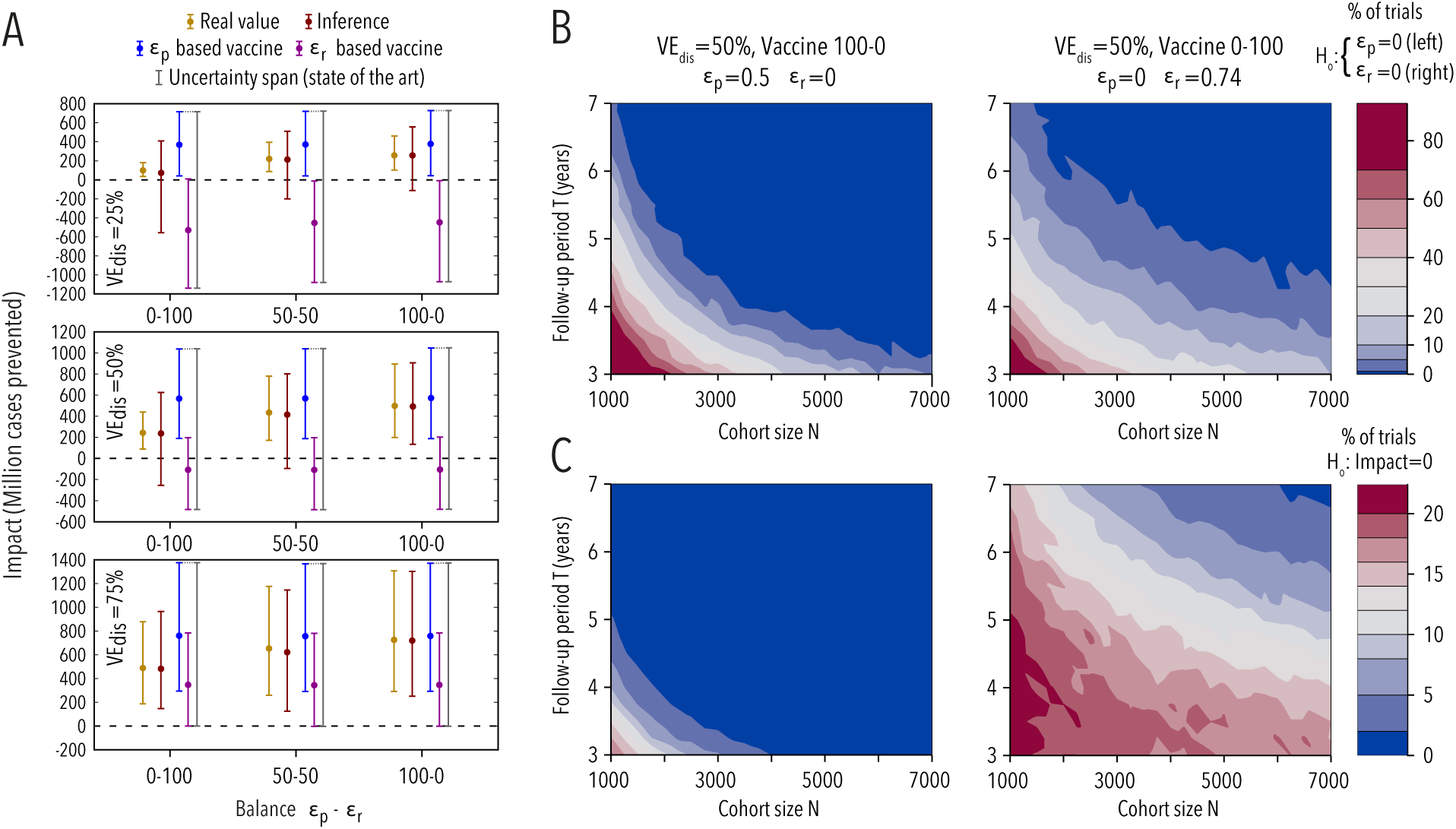
A. Real impact (i.e. derive from the real vaccine parameters), inferred impact (derived from the vaccine parameters obtained using our method) and state-of-the-art uncertainty for 9 different vaccines (3 different levels of efficacy at disease × 3 different balances between *ε_p_* and *ε_r_*). In all cases, the efficacy of the vaccine against infection is null (*ε_β_* = 0), while the vaccine characterization prior to impact estimations is made upon trials of *N* = 3000 and *T* = 4*y*. While the real impact (i.e. the impact foreseen from a vaccine that presents the real, underlying parameters *ε_β_*, *ε_p_*, *ε_r_*) carries only the uncertainty related to the spreading model; the inferred impact from our approach also involves the uncertainty given by the characterization of the vaccine. State-of-the-art uncertainty is obtained after simulating the impact with the two points of the curve (*ε_p_*; *ε_r_*) that will provide the most extreme impacts. B. Proportion of false negatives in the characterization of the vaccine (i.e. fraction of realizations that leads to a characterization of *ε_p_* (left) or *ε_r_* (right) that is not significantly larger than zero at a 95% CI). We study two different vaccines: *ε_p_*-based (left) and *ε_r_*-based (right), focusing on the characterization of the positive effect of the vaccine (i.e. *ε_p_* and *ε_r_* respectively). C. Probability of obtaining a positive impact for the two vaccines under study –*ε_p_*-based (left) and *ε_r_*-based (right)–. We assume that the impact obtained and its associated uncertainty correspond to a Normal Distribution with 95% CI.

To evaluate the advantages of our approach in this context, we compare the impact forecasts generated using the procedure described above to the most extreme impacts that can be obtained if, after measuring *V E_inf_* and *V E_dis_* from the synthetic trials, the efficacy against disease is mapped to all sort of possible combinations of values (ε_r_, *ε_p_*) that are compatible with it, a procedure that could beassimilated to the uncertainty provided by state of the art efficacy evaluation. As it can be seen in figure 3A, the degeneration between possible mechanisms underlying protection against disease translates into exacerbated levels of uncertainty that almost systematically prevent the rejection of the null hypothesis of null vaccine impact; an observation that is valid for a wide range of values of *V E*_dis_.

On the contrary, analyzing simulated trials’ data using our method substantially contributes to reduce the associated uncertainty of impact forecasts, in a factor ranging from 23% to 63% in the examples showed. The foreseen impacts showed in Figure 3A have two sources of uncertainties involved, those that come from the evaluation of vaccine parameters and those coming from the spreading model itself, except when we evaluate the real underlying vaccine for which we only have the latter. As shown in Figure 3A, in the case of vaccines that reducefast-progression, the inferred parameters of the vaccine introduce a significant increase inthe Confidence Interval of the forecasts. This situation is avoided in the case of vaccines that prevent fast-progression from taking place, whose confidence intervals are comparable with the uncertainty that is exclusive from the model.

Finally, we are interested in estimating the probability of registering a successful trial (i.e. a trial associated to a significant estimate in terms of vaccine descriptors and forecasted impact), as a function of both trial size and true, underlying vaccine features. To achieve this, we simulated clinical trials of different sizes and durations for the two extreme vaccines among those represented in figure 1E (both of which share the same reference level of protection *V E_dis_*(*T* = 4) = 50%), which act either by reducing proportion of fast progressors (*ε_r_* = 0, *ε_p_* = 0.50) or by slowing them down in their path to disease (*ε_r_* = 0.74, *ε_p_* = 0). In figure 3B, we show, as a function of trials dimensions, the probability of obtaininga valid trial simulation yielding an inferred value for the corresponding " statistically significant (95% CI not crossing zero). In figure 3C, we complement these results by characterizing how likely it is to obtain a clinical trial result associated to a forecast with significant impact.

## III. DISCUSSION

Despite TB being one of the diseases for which a vaccine was first developed, there is still much to learn about the mechanisms that TB vaccines, both BCG and novel candidates, unfold to disrupt the pathogen’s life cycle. Recent studies have challenged the classical view according to which BCG works solely by priming T-cell mediated adaptive immune responses, proposing more complex models of vaccine function. According to these new views, BCG vaccinationisalso able to boost host’s innate immune system, so providing an additional layer of non-specific, sustainable protection against infection, (usually referred to as “trained immunity”),which can be observed both in-vivo and in-vitro [28, 29]. How these observations might contribute to explain certain epidemiological aspects of BCG’s behavior is still unclear; like its ability to reduce *M.tuberculosis*infection risk [30]; its variable protection levels against TB disease [31], or their correlations with age, latitude and/or previous exposure to mycobacterial antigens [11, 12]. In any case, there is increasing scientific evidence suggesting that vaccines confer protection against infection and disease by mediating host-pathogen interactions in ways that,specially in the TB case, are more complex than classically accepted. An additional piece of this intricate puzzle, which we begin to analyze here, is that of understanding the possible repercussions of these improved vaccine descriptions on disease transmission dynamics.

In this work, we have described how vaccine protection against disease can manifest in different ways at the epidemiological level, e.g., by slowing down fast progression to disease(*ε_r_*-based vaccines) and/or preventing it completely (*ε_p_*-based vaccines), among others. As we have shown, these putative mechanisms of action cannot be disentangled by applying simple survival analysis. This limitation of standard methodologies when applied to TB is unexpectedly relevant, for we have shown that vaccines relying its protective action on these two mechanisms, even if they might appear as equally effective in the context of a clinical trial of vaccine efficacy, behave in remarkably different ways. As a result, this affects both the feasibility of their characterization as successful vaccines and the ultimate impact on TB burden reduction that they can provide. In this sense, *ε_r_*-based vaccines appear associated tolower prospected impacts for equivalent lectures of vaccine efficacy, and they are harder to analyze, as they would require bigger clinical trials of longer duration in order to be characterized at comparable levels of statistical significance. Therefore, distinguishing between these two different mechanisms of action improves early evaluation of TB vaccine candidates by contributing to a deeper description of vaccines’ expected behavior when applied on large populations and, more importantly, by avoiding previously unnoticed biases in their comparison, a crucial aspect in a context where resources are limited and committed only to the mostspromising candidates at each stage. As a result, considering the questions explored in thisstudy will contribute both to achieve a better interpretation of its outcomes as well as to improve the probability of success of a clinical trial of vaccine efficacy.

The method here proposed is instrumental in a number of situations, but has intrinsic limitations regarding both the dimensions of the trials that can be analyzed as well as the types of vaccines that can be characterized more accurately by using it. On the one hand, the framework proposed in this work is of application in the most favorable event of a vaccine being able to prevent TB infection (*ε_β_*-based vaccine) too. However, as explored in the appendix (see supplementary section S4), when protection against disease is primarily achieved through a reduction of the probability of infection, the considerations made in this work are quantitatively less relevant. In that case, the appearance of additional vaccine effects, either via *ε_p_* or via *ε_r_* yields more comparable vaccine behaviors, because the common, main vaccine effect on reducing the infection rate defines the impact expected for the vaccine.

On the other hand, in this work we have simulated trials of cohort sizes comprised between 1000 and 7000 individuals, during follow-up periods between 3 and 7 years. In the limit of short, small trials, low statistics constitutes a fundamental limitation that reduces the chances of obtaining a successful vaccine characterization, even for highly efficient vaccines. In turn, if the trial size and duration has to be increased, while increasing cohort size is always positive, implementing longer follow-ups may require different analytic approaches to those here presented. The reason is that the fundamental hypothesis behind our method is that all individuals progressing to disease during the trial can be associated to immediate progression upon infection: if the trial period is arbitrarily extended, such hypothesis does not hold anymore, and our method for estimating *ε_r_* and *ε_p_* would be biased.

In addition, our method of vaccine characterization can in principle be extended to more complex scenarios where more subtle phenomenologies can be considered. This includes to study the effect of vaccine heterogeneity, either within or across age (i.e. immunity waning), as well as to include in the simulated trials the dependence of some of the dynamical parameters on subjects’ age. Admittedly, even if such refinements might be relevant from a quantitative stand, other limitations regarding the amount of information that can be extracted from
a short clinical trial will most likely persist, regardless of these considerations. This comprises our intrinsic inability to estimate the effects of a vaccine on the slow dynamics ofLTBI from a short-course trial, as well as the existence of multiple unavoidable sources of uncertainty that affects the precision of model-based impact and cost-effectiveness evaluations, independently on how precise the efficacy of the vaccine itself is estimated from a trial.

Despite these limitations, the methodology presented in this work is instrumental for designing clinical trials that are more likely to succeed in the characterization of novel TB vaccines – and for providing a deeper characterization of them–, able to reduce uncertainty in impact forecasts evaluations.

## IV. METHODS

### A. Synthetic clinical trials simulations

To produce synthetic simulations of clinical trials, we first calibrated the baseline parameters of the transmission model to reflect the current epidemic situation in Worcester, South Africa, where the MVA85A study took place [18]. First, the transition rate from LTBI to disease is assumed to be *r_L_* = 7.5 × 10^−4^*y*^−1^, in accordance with previous bibliographical estimations [25]. The probability of fast-progression has been fixed to *p* = 0.375 which is compatible with previous observations about the remarkably high probability of developing fast-progression during the first months of life. The value used in this work is a reference to simulate trials conducted in newborns. Second, the baseline transmission rate was estimated to be *β* = 0.069*y*^−1^ to reproduce the proportion of infections in the control cohort of the MVA85A trial (12.8% after 2 years). Finally, the transition rate from fast latency to disease *r* = 0.97*y*^−1^ was calibrated to reproduce the same proportion of cases than in the control cohort of MVA85A trial (2.3% after 2 years), once all the other parameters are fixed. The fast-progression rate is also compatible with previous observations [25]. Thus, our approach is specific both to the site and age-strata of individuals joining the trial, which means that our inferred baseline parameters cannot be automatically used to simulate trials conducted in other sites, or age-cohorts, although similar re-calibrations are possible upon availability of reference data. Similarly, since BCG vaccination is mandatory in South Africa, the baseline parameters have embedded the eventual protective effects provided by the old vaccine.

Next, we arbitrarily define a vaccine by providing the triad of vaccine efficacies (*ε_β_*, *ε_r_*, *ε_p_*), describing its effects on the infection rate, the transition rate to disease, and the probability of fast progression, respectively. While a value of, for example, *ε_β_* = 0 means no protection against infection, and *ε_β_* = 1 means perfect protection, it is worth noticing that, in principle, vaccines where any of the three ε parameters is lower than zero are possible, as it is possible to observe a failed vaccine that indeed increases the risk of infection or disease with respect to the control cohort, as occurred in the MVA85A trial (although not significantly) [18].

Once all the dynamical parameters governing TB transmission dynamics in both cohorts are set up, we use an agent-based approach to simulate the evolution of *N* individuals per cohort during a follow-up period *T*. We use a discrete time step of 3 months to reproduce the temporal resolution between consecutive analyses (QFT for infection and/or TB diagnosis tests for active TB) in the MVA85A trial.

### B. Vaccine description (I): Estimation of *ε_r_*: truncated fit of transition rates from uncensored sub-cohorts

The well known differences between time-scales associated to LTBI and fast-progression to disease are captured, in terms of our elementary transmission model, by the relation *r_L_* ≪ *r*. This characteristic feature of TB transmission cycle allows us to assume that virtually all the individuals progressing to disease within the follow-up period of a clinical trial can be thought of as fast progressors. Furthermore, if we assume that transition from active disease upon infection is a Poisson process, -as it is done customarily in compartmental models in mathematical epidemiology [22, 25, 32]-, the theoretical probability distribution function (PDF) of the time *t* = *t*_dis._ − *t*_inf._ between infection and disease in the control cohort would correspond to an exponential curve *f*(*t*|*r*) = *re*^−*rt*^, from which the average transition time 〈*t*〉 = 1/*r* and its associated variance σ*_t_* = 〈*t*^2^〉 − 〈*t*〉^2^ = 1/*r*^2^ can be easily obtained by integrating the moments of the PDF.

However, in the practical context of a clinical trial, the period of measure cannot be arbitrarily extended, which implies that the maximum transition time that can be observed for a subject who was initially infected
at *t_inf_* is truncated at *t_max_* = *T* − *t_inf_*, where *T* stands for the follow-up period of the trial. This situation implies that the integrals needed to obtain the expected value of the transition time need to be truncated as well, which ultimately makes 〈*t*〉 to depend itself on *t_inf_*:

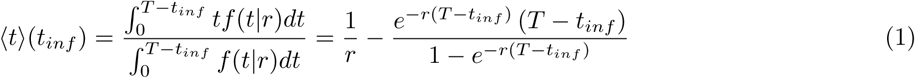

Similarly, by truncating the integrals of the second moment of the distribution we can obtain its dependence with time at infection, 〈*t*^2^〉(*t_inf_*)), and ultimately derive the corresponding expression for the variance of observed transition times as a function of *t_inf_*:

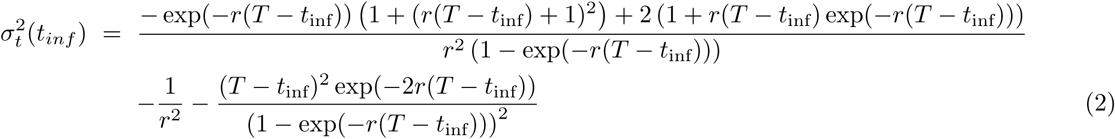

Equations 1 and 2 describe how observed transition times from infection to disease and their variance are expected to be biased towards lower values as the infections occur later during the trial; simply because the later the infection takes place, the less time available to observe a transition to disease is left. These expressions allow us to isolate the effect of that bias, and to infer, using only data from individuals developing active TB during the trial, the transition rate *r* within the control cohort, using a Maximum Likelihood approach (R package bbmle [33]) along with its confidence intervals (95% reported). Then, the exercise is repeated in the vaccine cohort, whose transition rate *r^v^*, in terms of our transmission model would be expressed as the product *r*(1 − *ε_r_*), which yields the following expression for the vaccine effect on fast progression rate *ε_r_*:

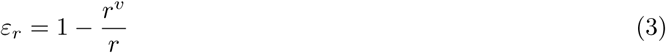

Finally, we obtain an estimation of the CI for *ε_r_* by propagating the independent uncertainties of *r^v^* and *r*.

### C. Vaccine description (II): estimation of *ε_β_* and *ε_p_*

The effect of the vaccine on the infection rate, codified in our model as *ε_β_*, coincides, by construction, with the vaccine efficacy against infection *V E_inf_*, and, as such, is inferred using Cox-regression (R package OIsurv [34]).

In order to infer the third vaccine effect -reduction of primo-infection probability *ε_p_*, we begin by describing the time evolution of the four sub-populations -*S*,*F*,*L* and *D*- of each cohort, making use of the following ordinary differential equations, mean field transmission model:

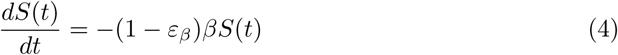

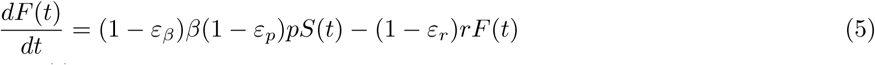

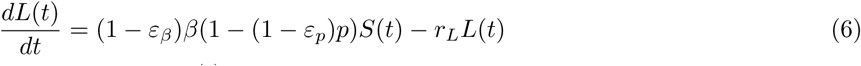

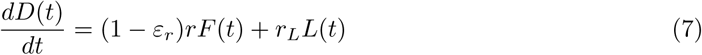

where the three vaccine descriptors (*ε_β_*, *ε_p_*, *ε_r_*) are absent (i.e. set to zero) in the control cohort. In this model we implicitly assume that the individuals in the cohorts correspond to a small fraction of the total population in the site, and thus, their contribution to overall transmission once sick can be neglected. By integrating this model analytically and independently in each cohort (see appendix, section S1), we can define the disease ratio *ρ* ≡ *D_v_*(*t*)/*D_c_*(*t*), and obtain an analytical expression for it that depends on the observation time *t*, and the vaccine descriptors (*ε_β_*; *ε_p_*; *ε_r_*):

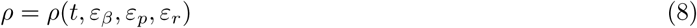

Then, we measure the vaccine reduction coefficient at the end of the trial *ρ*(*t* = *T*, *ε_β_*, *ε_p_*, *ε_r_*), and its uncertainty, which is propagated assuming that both *D_v_* and *D_c_* come from two independent binomial distributions (total number of tests equal to cohort size). then, using the independent estimators of *ε_β_* and *ε_r_* as well as their uncertainty estimates, obtained as detailed above, we get our final estimate of *ε_p_* and propagate its corresponding confidence interval.

In principle it is expected for the intrinsic efficacies of the vaccine to be comprised between 0 and 1, where 1 would mean a perfect efficacy and 0 no effect at all. However it is possible for a vaccine to have a negative effect. In the case of efficacies affecting rates (i.e. *ε_β_* and *ε_r_*) there is no formal lower limit and a rate equals to 1 (associated to ε = −∞) would mean an instantaneous process, although a conservative enough limit of -300 is implemented to avoid numerical instabilities. On the contrary, *ε_p_* works as a modifier of a probability, which implies that (1 − *ε_p_*)*p* has to be comprised between 0 and 1, introducing a lower limit for *ε_p_* that is 
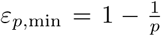
. Furthermore, the existence of such lower bound in *ε_p_* generates in turn an upper bound for *ε_r_*, since these two parameters are analytically bounded (see supplementary information, section S1) through a common value of the disease-ratio ρ = *D_v_*/*D_c_*. Notwithstanding this, the inference of *ε_r_* is agnostic to the value of *ρ* or *ε_p_*, and, as a consequence, for poor statistical settings –most often in the case of vaccines delaying fast-progression– some individual trial realizations lead to vaccine descriptor estimates that lie beyond these epidemiologically meaningful intervals for parameters *ε_p_* and *ε_r_*. Those ill-defined realizations, whose fraction is represented in figure 1E, are excluded from subsequent analyses.

### D. Obtention of global estimators and CI’s for (*ε_β_*, *ε_p_*, *ε_r_*) from simulated trials

In order to obtain global estimates and confidence intervals for vaccine descriptors we follow a three steps approach. First, we generate a set of 500 synthetic clinical trials for each vaccine analyzed. Second, for each of these simulated trials, we infer the values of the vaccine descriptors *ε_β_*, *ε_p_* and *ε_r_* along with their confidence intervals: that of *ε_β_* from Cox-regression, that of *ε_r_*, as explained in section IV B, and, finally, that of *ε_p_* propagated from the other two, and from the CI of the disease ratio ρ, as explained in the supplementary information (section S1). Finally, we assume that the true values of these parameters come from an unweighted mixture of normal distributions each of which is associated to the log transform of one minus the outcome of each simulated trial. The final value and CI of each of the three vaccine descriptors is associated to the median and 95% CI of such distribution mixture, back in the linear scale. Through this approach we get a global estimation of the accuracy and precision of our method as a function on the predefined vaccine’s characteristics and trial dimensions, which we have introduced in the TB spreading model for the forecasts of vaccine impacts (and their correspondent Confidence Interval).

### E. Model based impact evaluations of TB vaccines

We have used the estimates of the efficacy and their respective CIs (as described in the previous section) to evaluate and compare the potential impact of these hypothetical vaccines when applied on larger populations. To do so, we take advantage of the detailed TB transmission model described in [21], developed by the authors as a tool for the impact estimation of novel TB vaccines. This model is a generalization of increased complexity of the reduced transmission model sketched in figure 1A and formalized around the simple ODE system showed through equations 4-7. The most important difference between both formulations is that, while the elementary transmission model described in this work is suited to capture the time evolution of the fraction of susceptible, infected and sick individuals within the trials’ cohorts (constituted by infants) during the development of the study, the more complex version developed in [21] was designed to produce impact evaluations of novel vaccines in broad epidemiological settings, and, as such, it describes how do these proportions of susceptible, infected, and sick individuals evolve in entire age-structured populations spanning different countries during more prolonged periods of time.

The technical specifications of this detailed age-structured model of TB transmission can be found in [21] and supplementary information (Section S3). It contemplates the existence of two cohorts of individuals -vaccinated and non-vaccinated-, two paths to disease -fast and slow- and six different situations of disease, depending on treatment status (present or absent) and on its aetiology -pulmonary (smear positive/negative) vs. non pulmonary-. Regarding treatment results, the model explicitly describes the main outcomes defined by WHO data schemes: treatment completion, default, failure and death, as well as natural recovery. Furthermore, several types of infection events are taken into account, including infection of previously unexposed individuals, exogenous reinfections of previously infected subjects, and mother-child transmission.

For the sake of the impact evaluations presented in this work, the model is calibrated in Ethiopia. Using demographic data from the UN population division database [35] as well as aggregated burden estimates reported for these countries in the interval [36], we calibrated the model to reproduce the TB epidemics outlook in this region between 2000 and 2015, as detailed in [21]. Finally, we evaluate the putative impact that the different vaccines characterized in this study would produce in that region, if applied in 2025 through an immunization campaign targeting newborns, as the number of active TB cases prevented in the 25 years following the implementation of the campaign (2025-2050).

## References

[1] WHO. Global Tuberculosis Report 2017. World Health Organization, 2017.

[2] Christopher Au-Yeung, Steve Kanters, Erin Ding, Philippe Glaziou, Aranka Anema, Curtis L Cooper, JS Montaner, Robert S Hogg, and Edward J Mills. Tuberculosis mortality in hiv-infected individuals: a cross-national systematic assessment. Clin Epidemiol, 3:21–29, 2011.

[3] Cristiana J. Silva and Delfim F. M. Torres. A tb-hiv/aids coinfection model and optimal control treatment. Discrete and Continuous Dynamical Systems, 35(9):4639–4663, 2015.

[4] Amita Jain and Pratima Dixit. Multidrug resistant to extensively drug resistant tuberculosis: what is next? Journal of biosciences, 33(4):605–616, 2008.

[5] James M Trauer, Justin T Denholm, and Emma S McBryde. Construction of a mathematical model for tuberculosis transmission in highly endemic regions of the asia-pacific. Journal of theoretical biology, 358:74–84, 2014.

[6] Laura C Rodrigues, Vinod K Diwan, and Jeremy G Wheeler. Protective effect of bcg against tuberculous meningitis and miliary tuberculosis: a meta-analysis. International journal of epidemiology, 22(6):1154–1158, 1993.

[7] Graham A Colditz, Timothy F Brewer, Catherine S Berkey, Mary E Wilson, Elisabeth Burdick, Harvey V Fineberg, and Frederick Mosteller. Efficacy of bcg vaccine in the prevention of tuberculosis: meta-analysis of the published literature. Jama, 271(9):698–702, 1994.

[8] Paul EM Fine. Variation in protection by bcg: implications of and for heterologous immunity. The Lancet, 346(8986):1339–1345, 1995.

[9] B Bourdin Trunz, PEM Fine, and C Dye. Effect of bcg vaccination on childhood tuberculous meningitis and miliary tuberculosis worldwide: a meta-analysis and assessment of cost-effectiveness. The Lancet, 367(9517):1173–1180, 2006.

[10] Punam Mangtani, Ibrahim Abubakar, Cono Ariti, Rebecca Beynon, Laura Pimpin, Paul EM Fine, Laura C Rodrigues, Peter G Smith, Marc Lipman, Penny F Whiting, et al. Protection by bcg vaccine against tuberculosis: a systematic review of randomized controlled trials. Clinical infectious diseases, 58(4):470–480, 2014.

[11] Mauricio L Barreto, Daniel Pilger, Susan M Pereira, Bernd Genser, Alvaro A Cruz, Sergio S Cunha, Clemax Sant’Anna, Miguel A Hijjar, Maria Y Ichihara, and Laura C Rodrigues. Causes of variation in bcg vaccine efficacy: Examining evidence from the bcg revac cluster randomized trial to explore the masking and the blocking hypotheses. Vaccine, 32(30):3759–3764, 2014.

[12] Sergio Arregui, Joaquín Sanz, Dessislava Marinova, Carlos Martín, and Yamir Moreno. On the impact of masking and blocking hypotheses for measuring the efficacy of new tuberculosis vaccines. PeerJ, 4:e1513, February 2016.

[13] WHO. Global Tuberculosis Report 2015. World Health Organization, 2015.

[14] Dessislava Marinova, Jesus Gonzalo-Asensio, Nacho Aguilo, and Carlos Martin. Recent developments in tuberculosis vaccines. Expert review of vaccines, 12(12):1431–1448, 2013.

[15] Ann M Ginsberg, Morten Ruhwald, Helen Mearns, and Helen McShane. Tb vaccines in clinical development. Tuberculosis, 99:S16–S20, 2016.

[16] Helen A Fletcher. Correlates of immune protection from tuberculosis. Current molecular medicine, 7(3):319–325, 2007.

[17] Kamlesh Bhatt, Sheetal Verma, Jerrold J Ellner, and Padmini Salgame. Quest for correlates of protection against tuberculosis. Clinical and Vaccine Immunology, 22(3):258–266, 2015.

[18] Michele D Tameris, Mark Hatherill, Bernard S Landry, Thomas J Scriba, Margaret Ann Snowden, Stephen Lockhart, Jacqueline E Shea, J Bruce McClain, Gregory D Hussey, Willem A Hanekom, et al. Safety and efficacy of mva85a, a new tuberculosis vaccine, in infants previously vaccinated with bcg: a randomised, placebo-controlled phase 2b trial. The Lancet, 381(9871):1021–1028, 2013.

[19] Anne O’Garra, Paul S Redford, Finlay W McNab, Chloe I Bloom, Robert J Wilkinson, and Matthew PR Berry. The immune response in tuberculosis. Annual review of immunology, 31:475–527, 2013.

[20] M Elizabeth Halloran, Kari Auranen, Sarah Baird, Nicole E Basta, Steve Bellan, Ron Brookmeyer, Ben Cooper, Victor DeGruttola, James Hughes, Justin Lessler, et al. Simulations for designing and interpreting intervention trials in infectious diseases. bioRxiv, page 198051, 2017.

[21] Sergio Arregui, Joaquin Sanz, Dessislava Marinova, Maria Jose Iglesias, Sofia Samper, Carlos Martin, and Yamir Moreno. A data-driven model for the assessment of age-dependent patterns of tuberculosis burden and impact evaluation of novel vaccines. bioRxiv, page 112409, 2017.

[22] Christopher Dye, Geoffrey P Garnett, Karen Sleeman, and Brian G Williams. Prospects for worldwide tuberculosis control under the {WHO} {DOTS} strategy. The Lancet, 352(9144):1886–1891, 1998.

[23] T Lietman and SM Blower. Potential impact of tuberculosis vaccines as epidemic control agents. Clinical Infectious Diseases, 30(Supplement 3):S316–S322, 2000.

[24] Elad Ziv, Charles L Daley, and Sally Blower. Potential public health impact of new tuberculosis vaccines. Emerg Infect Dis, 10(9):1529–1535, 2004.

[25] Laith J Abu-Raddad, Lorenzo Sabatelli, Jerusha T Achterberg, Jonathan D Sugimoto, Ira M Longini, Christopher Dye, and M Elizabeth Halloran. Epidemiological benefits of more-effective tuberculosis vaccines, drugs, and diagnostics. Proceedings of the National Academy of Sciences, vol (33):13980–13985, 2009.

[26] Gwenan M Knight, Ulla K Griffiths, Tom Sumner, Yoko V Laurence, Adrian Gheorghe, Anna Vassall, Philippe Glaziou, and Richard G White. Impact and cost-effectiveness of new tuberculosis vaccines in low-and middle-income countries. Proceedings of the National Academy of Sciences, 111(43):15520–15525, 2014.

[27] Dany Pascal Moualeu, Martin Weiser, Rainald Ehrig, and Peter Deuflhard. Optimal control for a tuberculosis model with undetected cases in cameroon. Communications in Nonlinear Science and Numerical Simulation, 20(3):986–1003, 2015.

[28] Johanneke Kleinnijenhuis, Jessica Quintin, Frank Preijers, Leo AB Joosten, Daniela C Ifrim, Sadia Saeed, Cor Jacobs, Joke van Loenhout, Dirk de Jong, Hendrik G Stunnenberg, et al. Bacille calmette-guerin induces nod2- dependent nonspecific protection from reinfection via epigenetic reprogramming of monocytes. Proceedings of the National Academy of Sciences, 109(43):17537–17542, 2012.

[29] Eva Kaufmann, Joaquin Sanz, Jonathan L. Dunn, Laura S. Mendonca, Pacis Alain, Fanny Tzepelis, Erwan Pernet, Anne Dumaine, Florence Mailhot-Léonard, Eisha Ahmed, Jad Belle, Rickvinder Besla, Bruce Mazer, Irah L. King, Anastasia Nijnik, Barreiro Luis B., and Maziar. Divangahi. Bcg educates hematopoietic stem cells to generate protective innate immunity against tuberculosis. Cell, 172(1):XX–XX, 2018.

[30] Ahmet Soysal, Kerry A Millington, Mustafa Bakir, Davinder Dosanjh, Yasemin Aslan, Jonathan J Deeks, Serpil Efe, Imogen Staveley, Katie Ewer, and Ajit Lalvani. Effect of bcg vaccination on risk of mycobacterium tuberculosis infection in children with household tuberculosis contact: a prospective community-based study. The Lancet, 366(9495):1443–1451, 2005.

[31] A Roy, M Eisenhut, RJ Harris, LC Rodrigues, S Sridhar, S Habermann, L Snell, P Mangtani, I Adetifa, A Lalvani, et al. Effect of bcg vaccination against mycobacterium tuberculosis infection in children: systematic review and meta-analysis. Bmj, 349:g4643, 2014.

[32] Herbert W Hethcote. The mathematics of infectious diseases. SIAM review, 42(4):599–653, 2000.

[33] Ben Bolker et al. bbmle: Tools for general maximum likelihood estimation, 2010.

[34] David Diez. Survival analysis in r. OpenIntro. org, 2013.

[35] UN. Population division database. http://esa.un.org/unpd/wpp/index.htm, (accessed November 2016).

[36] WHO. Tuberculosis database. http://www.who.int/tb/country/en/index.html, (accesed November 2016).

